# Pyrroloquinoline Quinone inhibits PCSK9, augments LDLR recycling and enhances the effect of atorvastatin

**DOI:** 10.1101/2022.03.20.485060

**Authors:** Karan Devasani, Anuradha Majumdar, Dimple Jhonsa, Rachna Kaul

## Abstract

High fat diet intake or statin therapy are known to increase the expression of PCSK9, a key factor involved in the lipid metabolism. Inhibition of PCSK9 is a potential novel strategy for treatment of CVD. Aim of the present study is to investigate the effect of PQQ and atorvastatin on PCSK9 expression in high fat, 10% fructose diet (HFFD) fed rats. Rats received HFFD for 10 weeks followed by treatment of 5 weeks with ATS 10 or 20 mg/kg, PQQ 10 or 20 mg/kg, *p.o. per se* or in combinations. The effect of the treatments on serum lipid profile, transaminases and PCSK9 concentrations was estimated. Hepatic expressions of PCSK9 and LDLR were quantified by real time quantitative Polymerase chain reaction (RT-qPCR) and western blotting analysis. Compared with the positive control, rats receiving PQQ and ATS showed significant decrease in serum levels of triglycerides, total cholesterol, LDL, VLDL, non-HDL and atherogenic index. Further, drug treatment resulted in decreased serum ALT and AST levels (*P* < 0.0001). PQQ in combination with atorvastatin reduced serum PCSK9 levels (*P* < 0.0001) while downregulating the hepatic expression of PCSK9 (*P* < 0.0001) and increasing the expression of LDL-R (*P* < 0.001) and rescuing its degradation. PQQ along with atorvastatin prevented the LDLR degradation that was induced in the HFFD fed rats by decreasing PCSK9 expression. The results support an improvement in cholesterol management by adding PQQ to statin treatment.

**Graphical abstract:** 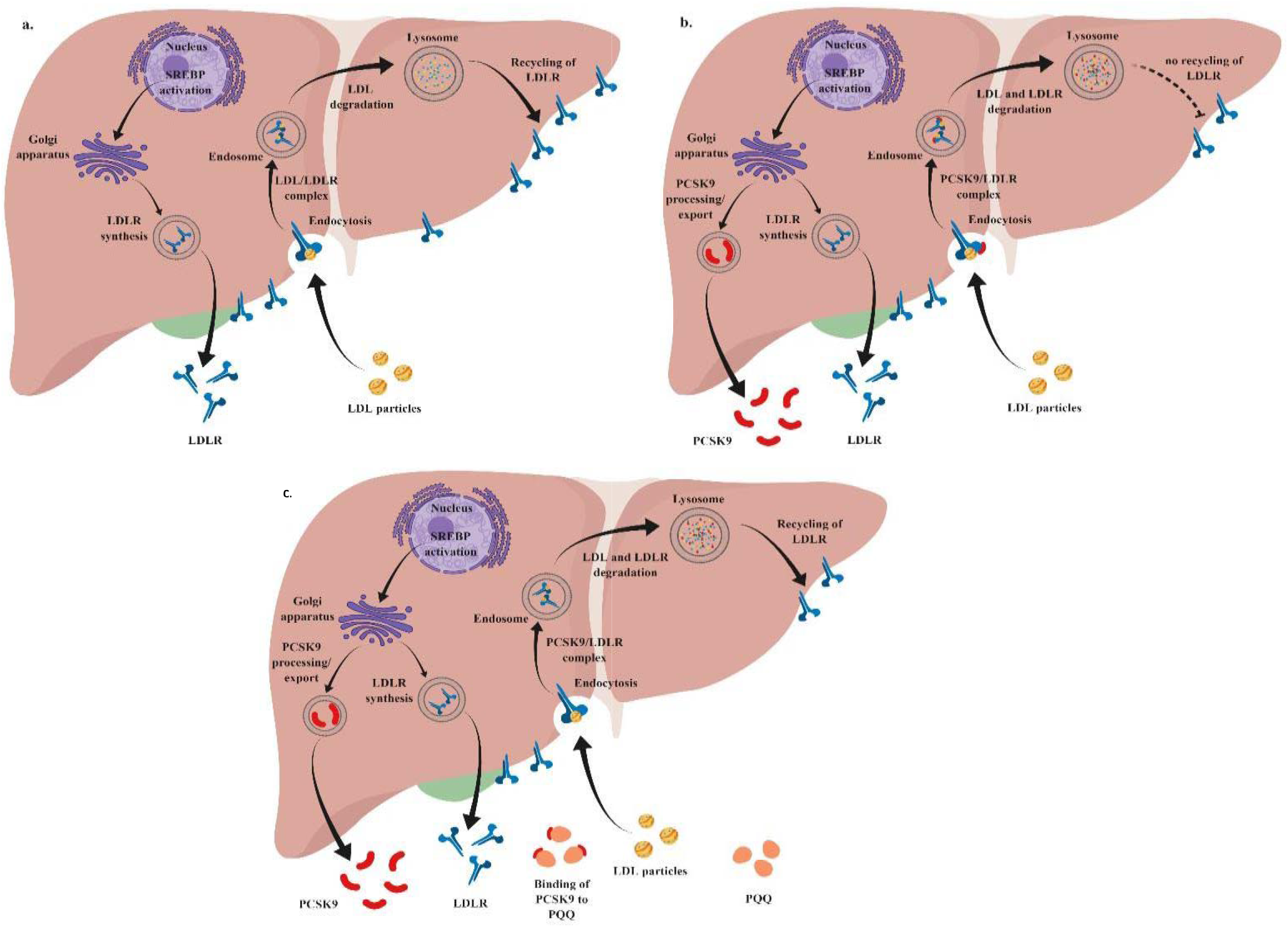

## Introduction

Hypercholesterolemia and obesity have become a pandemic health problem and is associated with clusters of diseases such as hypertension, atherosclerosis, insulin resistance and Type 2 Diabetes. It is well established that LDL and HDL are among the important predictors of cardiovascular events. LDL is indicated as an independent and most potent modifiable risk factor for originating cardiovascular diseases [1]. Proprotein convertase subtilisin/kexin type 9 (PCSK9), a serine protease involved in LDL metabolism is mainly expressed in the liver, intestine and kidney. It maintains the plasma LDL-C levels by regulating the hepatic cell surface LDL receptor (LDLR) to lysosomal degradation. Gain-of-function mutations of the PCSK9 is associated with hypercholesterolemia and increased risk of cardiovascular events [2] [3]. Thus, PCSK9 inhibits normal recycling of LDLR to the cell surface results in reduced density of LDLR and consequently increased accumulation LDL-C in the circulation [4]. Conversely, loss-of-function mutations are linked to low plasma LDL-C levels and a reduction of cardiovascular risk without known unwanted effects on individual health [5]. Statins such as Lipitor®, Zocor^®^ and crestor® are the most widely prescribed medications for lowering blood cholesterol that have been touted as the best way to manage cholesterol [6]. It is estimated that 20% of individuals show some degree of intolerance towards statin therapy and within 1-2 years after initiation of therapy, 40-75% of patients discontinued [7]. A systematic review gauged that 1.9 million patients consuming high doses of statin therapy have poor glycemic control with increased risk of new onset of diabetes mellitus in nondiabetics [8]. Millions of patients are prescribed statins where they really don’t need it. Statins are notorious for eliciting muscle pain, mitochondrial damage, Type 2 Diabetes, chronic low-grade inflammation (metainflammation), neurological problems and sexual dysfunction.

A meta-analysis of numerous randomized trials has established that statins delays cardiovascular events and prolongs survival by reducing LDL-C [9]. This reduction in LDLC is by thwarting the rate-limiting step in cholesterol synthesis, upregulating the expression of hepatic LDLR and increased clearance of LDL-C from circulation [10]. However, statins are known to not only upregulate LDL receptors, but also simultaneously upregulate expression and secretion of PCSK9 *via* a pathway involving the sterol regulatory elementbinding protein-2 (SREBP-2). PCSK9 decreases the number of LDLR in hepatocytes and results in increased circulation of LDL-C [11]. These contrary outcomes of statin therapy results in suboptimal or failure in therapeutic outcomes in hyperlipidemia and more seriously leads to statin intolerance.

Pyrroloquinoline quinone is an anionic water-soluble compound that serves as a redox cofactor and is ubiquitous in nature. In both plants and animals, it acts as a novel growth factor. It plays a central role in the regulation of cellular energy metabolism and mitochondrial biogenesis [12]’ [13]. It is suggested that PQQ have a role in stimulating immunity, protecting neurons and has antioxidant, anti□inflammatory and anticancer properties [14] [15] [16], which are mediated by a decrease in the release of inflammatory factors thus attenuating inflammatory diseases [17]. It has been reported that PQQ reduces myocardial infarct size and improves cardiac function and can protect against several types of oxidative damages and irradiation injury and exert an effective cardioprotective effect [16]’ [18]. However, the role of PQQ *per se* or its therapeutic potential along with statins has not been evaluated in hypercholesterolemia or obesity. Moreover, its impact on the PCSK9-LDLR pathway remains to be elucidated.

Therefore, the current study aimed to evaluate the effect of PQQ supplementation with atorvastatin in a HFFD model in rats with emphasis on exploring the impact on serum lipid profile and PCSK9 along with hepatic expression of PCSK9 and LDLR, essentially to understand role in lipid homeostasis.

## Materials and methods

Male Sprague Dawley rats, weighing around 180-200 g were housed in polypropylene cages, acclimatized to laboratory conditions maintained at room temperature 25±2°C, humidity 65±5%, 12-hours light-dark cycle and had free access to water and standard pelleted laboratory animal diet. After one week of acclimatization, rats received High Fat, 10% Fructose Diet (HFFD) (4,935 kcal/kg; 56.04% calories from carbohydrate, 12.36% calories from proteins, 24.44% calories from fats, 4.70% calories from dietary fiber, 10% fructose solution in fresh water) to induce obese hypercholesteremia rat model [17] [19] for 10 weeks and randomly assigned into eight groups (n=6).

Rats in Normal control group (Group I) were fed standard rat chow (3,020 kcal/kg; 52% calories from carbohydrate, 16.65% calories from proteins, 3.72% calories from fats, 5.5% calories from dietary fiber) and freshwater *ad libitum* throughout the study. Group II is considered as positive control (HFFD), groups from III to VIII received HFFD were treated with two dosages of PQQ (10 mg/kg or 20 mg/kg,*p.o*.), alone or in combination with atorvastatin (10 mg/kg or 20 mg/kg, *p.o*.) for 5 weeks [20]. Dose selection for ATS was done based on the earlier reports [21] [22] and for PQQ as mentioned in GRAS notice of PQQ [23]. At the end of the experiment, the blood samples were collected in EDTA-coated tubes and all animals were euthanized by CO_2_ inhalation. Livers were isolated to assess hepatic LDLR and PCSK9 protein levels and stored at −80°C until analysis. Animal experiments were reviewed and approved by the Institutional Animal Ethics Committee (IAEC), protocol no. CPCSEA-BCP/2016-02/12 and were in accordance with the guidelines of the Committee for the Purpose of Control and Supervision of Experiments on Animals (CPCSEA), Government of India.

### Serum Lipids, Transaminase and PCSK9 levels

Serum triglyceride, total cholesterol, low density lipoprotein cholesterol (LDL-C), Very low-density lipoprotein cholesterol (VLDL-C), high density lipoprotein cholesterol (HDL-C), non-HDL-C, Alanine transaminase (ALT), Aspartate transaminase (AST) was determined using commercial diagnostic kits (Erba Diagnostics, Mannheim, Germany). The concentrations of serum and hepatic PCSK9 levels were determined using ELISA kit according to manufacturer’s protocol (Elabscience®, Houston, Texas).

### Hepatic PCSK9 and LDLR gene and protein levels

#### mRNA expression of PCSK9 and LDLR

Total RNA was isolated using the PureLink® RNA mini kit (Invitrogen,) as per the manufacturer’s guidelines. Total RNA isolated was analyzed using Qubit® 3.0 Fluorometer (Invitrogen). Relative quantifications of PCSK9 and LDLR were carried out using Verso SYBR® Green 1-Step qRT-PCR ROX Mix (Thermo Scientific). Specific primers were synthesized commercially (Integrated DNA Technologies, Leuven, Belgium), with the following sequences: PCSK9: 5’-ATTGTACCTGTGGCTGGACG-3’ (forward); 5’-CCAGGATGAGACGCCAAGTT-3’ (reverse); LDLR: 5’-TTTGCAGCGGGAACATTTCG-3’ (forward); 5’-CACAGCTGAACTCACCGGAT-3’ (reverse); β-Actin: 5’-AGGCTGTGTTGTCCCTGTATG-3’ (forward); 5’-AACCGCTCATTGCCGATAGT-3’ (reverse)

The polymerase chain reaction condition for amplification are as follows: 95°C for 15 min for thermostat activation, followed by 40 cycles of denaturation at 95°C for 15 s, and combined annealing and extension at 50-55°C for 30 s and 72°C for 30 s respectively. The mRNA levels of PCSK9 and LDLR were normalized with the values of β-Actin. Results are expressed as fold changes of threshold cycle (Ct) value compared with controls using the 2^(-ΔΔCt)^ method [24]

#### Western blot analysis

Liver tissue lysates were homogenized in ice cold RIPA buffer containing the protease cocktail inhibitor as described previously [25]. Total protein concentrations were quantified by a BCA protein assay kit according to manufacturer’s protocol (Pierce BCA protein assay kit, catalogue no.: 23225). The protein samples electrophoresed by SDS-PAGE were electroblotted on PVDF by applying a current of 0.8 mA/h in a wet-transfer unit. Following transfer, the membrane was blocked with 3% BSA in TBS for 2 h at room temperature and incubated with the desired primary antibody overnight at 4°C. Following dilutions of antibodies were used: anti-PCSK9, 1:1000 (abclonal cat. no.: A7860), anti-LDLR, 1:600 (Santa Cruz Biotechnology, cat. no.: SC-8823), anti-β-Actin, 1:1000 (abclonal cat. no.: AC006). Post washing with TBST 3X the bound antibody complexes were detected using enhanced chemiluminescence (ECL) system (Pierce™ ECL western blotting substrate kit, catalogue no: 32106). Band intensities were quantified with Image J software and normalized to β-actin [20].

### Statistical analyses

Data analyses were performed using GraphPad Prism 7 software (GraphPad Software, San Diego, CA, USA) with results expressed as mean ± SEM. The significant differences among the experimental groups were compared by one-way analysis of variance (ANOVA), followed by Dunnett’s multiple comparison test as a *post-hoc* analysis. *P* values < 0.05 were considered nominally significant when compared with positive control and normal control groups.

## Results

### Effect of PQQ, atorvastatin and combination on serum lipid profile, transaminases and PCSK9 levels

Compared to positive control group, PQQ (10 mg/kg or 20 mg/kg) treatment decreased average serum TC (56 ± 3.53 mg/dL, *P* < 0.0001; 54.75 ± 3.81 mg/dL, *P* < 0.0001), TG (87.4 ± 6.56 mg/dL, *P* < 0.001; 72.8 ± 8.33 mg/dL, *P* < 0.0001), LDL (22.15 ± 1.45 mg/dL, *P* < 0.0001; 19.56 ± 2.01 mg/dL, *P* < 0.0001), VLDL (15.65 ± 0.82 mg/dL, *P* < 0.0001; 15.32 ± 0.92 mg/dL, *P* < 0.0001), non-HDL (56 ± 3.536 mg/dL, *P* < 0.0001; 54.75 ± 3.81 mg/dL, *P* < 0.0001). While in combination with atorvastatin (ATS+PQQ 20 mg/kg) the figures improved further with decreased TC (54.2 ± 1.46 mg/dL, *P* < 0.0001), LDL(14.95 ± 1.72 mg/dL, *P* < 0.0001), VLDL (13.28 ± 0.67 mg/dL, *P* < 0.0001), non-HDL (54.2 ± 1.46, mg/dL, *P* < 0.0001). Compared with positive control and atorvastatin treated groups, the combination treated groups revealed increased HDL (29 ± 1.73 mg/kg, *P* < 0.01; 31.4 ± 1.28 mg/kg, *P* < 0.0001) while TC, TG, LDL, VLDL reductions after treatments with atorvastatin, PQQ, and their combination were confined to non-HDLs (Figure 1a-f).

**Figure 1:**
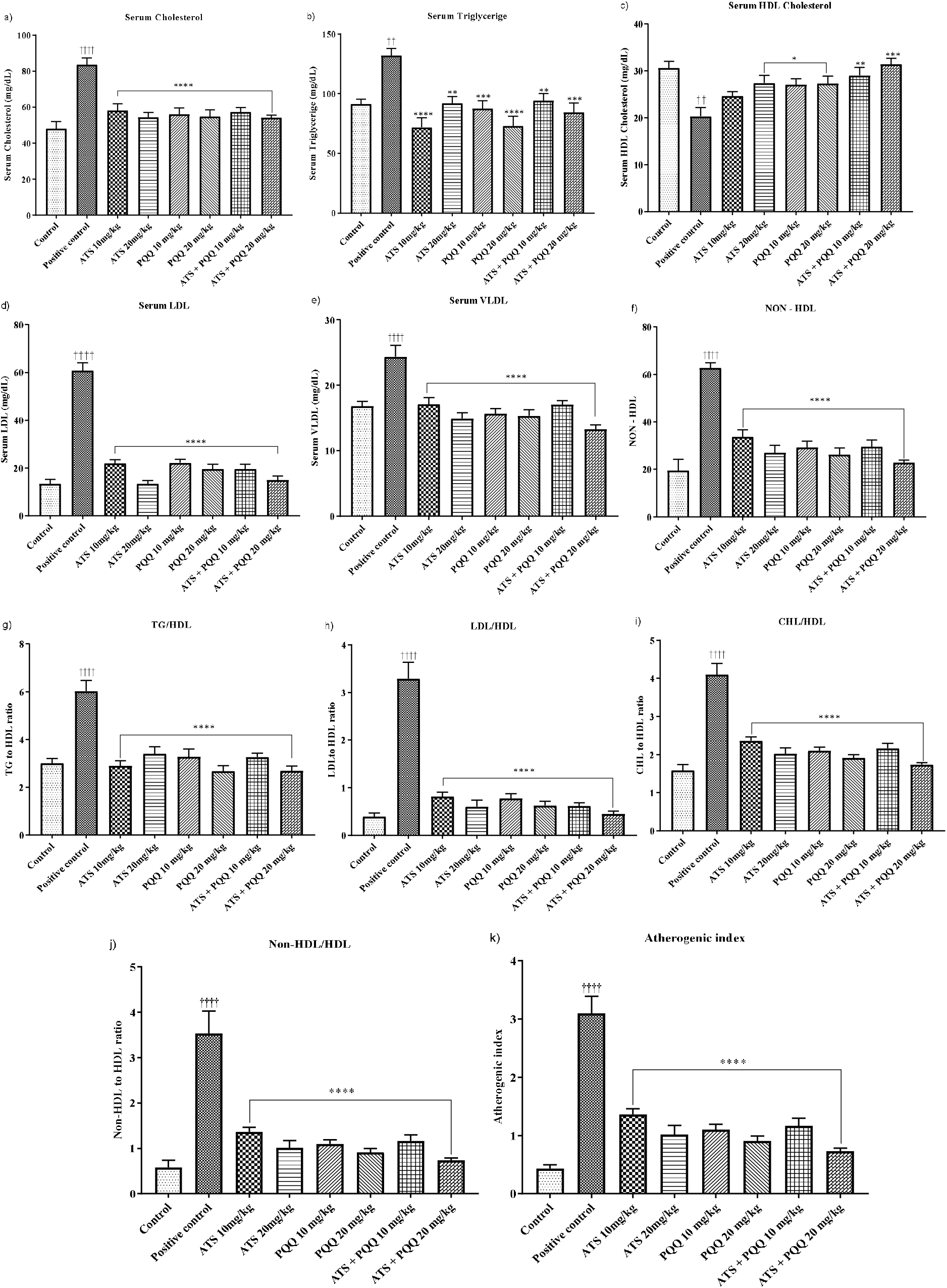
Effect of PQQ and ATS on serum lipid profile in HFFD fed male rats. a) Serum cholesterol, b) Serum Triglyceride, c) Serum HDL cholesterol, d) Serum LDL, e) Serum VLDL, f) Serum Non-HDL, g) TG/HDL, h) LDL/HDL, i) CHL/HDL, j) non-HDL/HDL, k) Atherogenic index. Data are means± SEM. ^††††^*P* < 0.0001; ^†††^*P* < 0.001; ^††^*P* < 0.01 when compared to the control group. \*\*\*\**P* < 0.0001; \*\*\**P* < 0.001; \*\**P* < 0.01; \**P* < 0.05 when compared with positive control group. Statistical significance was determined by one-way ANOVA with Dunnett’s post-hoc test.

Lipoprotein ratios and atherogenic index were calculated to predict cardiovascular risk factors. In comparison to normal control, HFFD fed positive control rats showed significant increase in the ratios of TG/HDL (6.022 ± 0.4519, *P* < 0.0001), LDL/HDL (3.29 ± 0.34, *P* < 0.0001), CHL/HDL (4.095 ± 0.29, *P* < 0.0001), non-HDL/HDL (3.532 ± 0.49, *P* < 0.0001) and atherogenic index (3.095 ± 0.29, *P* < 0.0001). The ratios and atherogenic indices were observed to significantly reduce indicating lowered cardiovascular risk with treatments of atorvastatin, PQQ *per* se and their combinations (Figure 1g-k).

### Effect of PQQ, atorvastatin and combination on serum transaminases and PCSK9 levels

Consumption of HFFD augured significant increase in serum circulating levels of ALT (831.2 ± 24.13 mg/dL, *P* < 0.0001) and AST (250.8 ± 6.82 mg/dL, *P* < 0.0001) in the positive control rats when compared with normal control. These perturbations of the liver function markers were restored to normal levels by treatment with atorvastatin, PQQ *per* se and their combination (Figure 2a-b).

**Figure 2:**
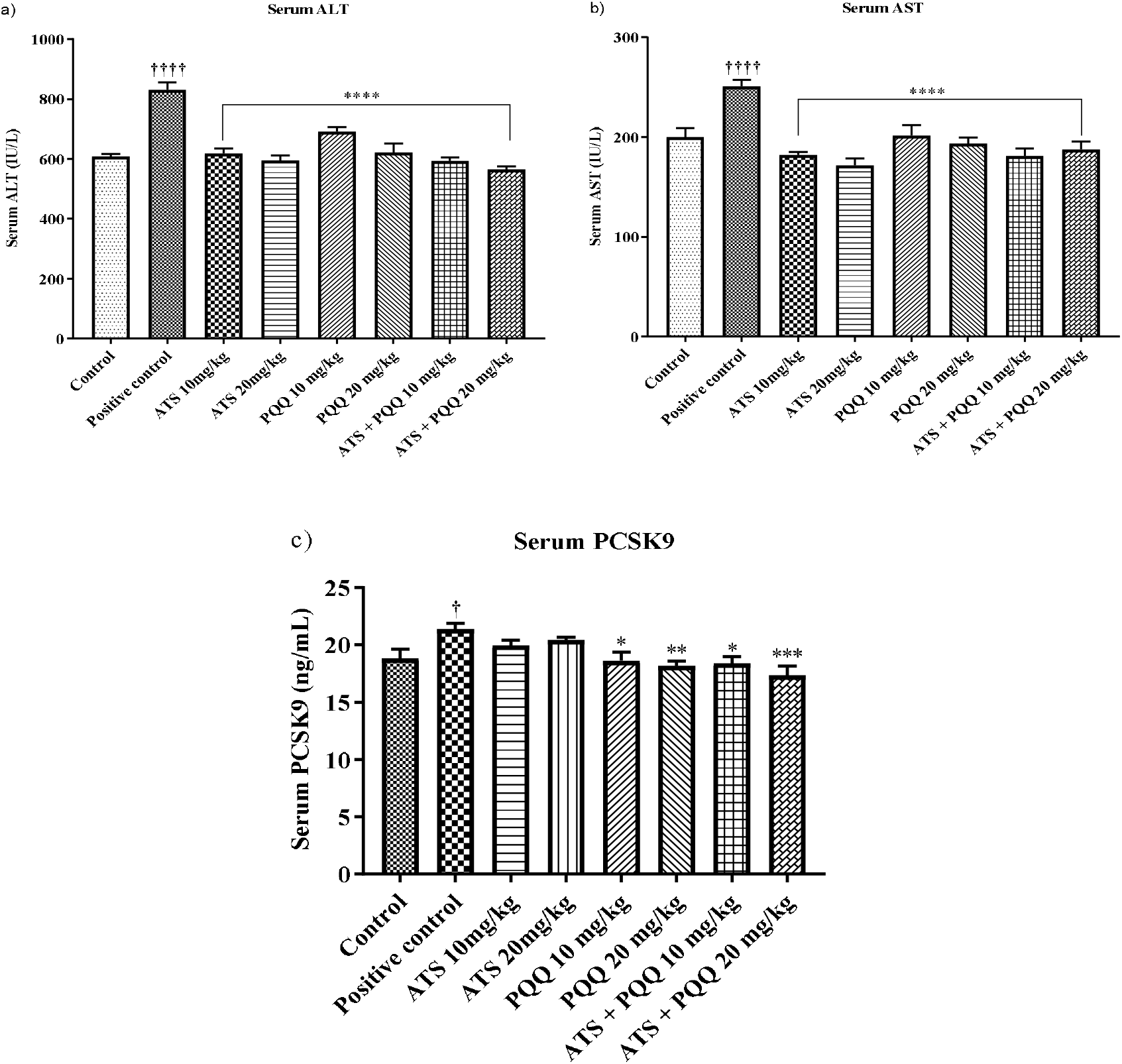
Effect of PQQ and ATS on serum ALT and AST in HFFD fed male rats. a) Serum ALT, b) Serum AST and c) serum PCSK9. Data are means ± SEM. ^††††^*P* < 0.0001 when compared to the control group. \*\*\*\**P* < 0.0001; \*\*\**P* < 0.001; \*\**P* < 0.01; \**P* < 0.05 when compared with positive control group. Statistical significance was determined by one-way ANOVA with Dunnett’s post-hoc test.

Serum PCSK9 levels were significantly higher (21.38 ± 0.51 ng/mL, *P* < 0.05) in rats fed with HFFD compare to normal control group (18.79 ± 0.81 ng/mL) demonstrating that HFFD could upregulate the hepatic expression of PCSK9 and its release in blood. Compared with positive control group, the rats treated with atorvastatin *per se*, PQQ 20 mg/kg and the combination (ATS+PQQ 20mg/kg) revealed significant noteworthy lower levels of serum circulating PCSK9 (18.18 ± 0.40 ng/mL, *P* < 0.01; 17.33 ± 0.83 ng/mL, *P* < 0.001) (Figure 2c).

### Effect of PQQ, atorvastatin and combination on mRNA and protein expression of hepatic PCSK9 and LDLR

Treatment with PQQ 10 or 20 mg/kg *per se* (0.39 ± 0.10, *P* < 0.0001; 0.20 ± 0.01, *P* < 0.0001) and in combination with atorvastatin (0.35 ± 0.07, *P* < 0.0001; 0.19 ± 0.02, *P* < 0.0001) for 5 weeks significantly decreased the hepatic PCSK9 expression in rats fed with HFFD as compared to positive control. This decrease in the levels of PCSK9 impacted by upregulating the mRNA expression of LDLR in the PQQ 20 mg/kg *per se* (3.32 ± 0.61, *P* < 0.01) and the combination (3.52 ± 0.40, *P* < 0.01; 3.95 ± 0.39, *P* < 0.001) treated rats when compared to positive control and the group treated with atorvastatin 10 or 20 mg/kg (1.72 ± 0.55; 1.46 ± 0.19) (Figure 3a-b).

**Figure 3:**
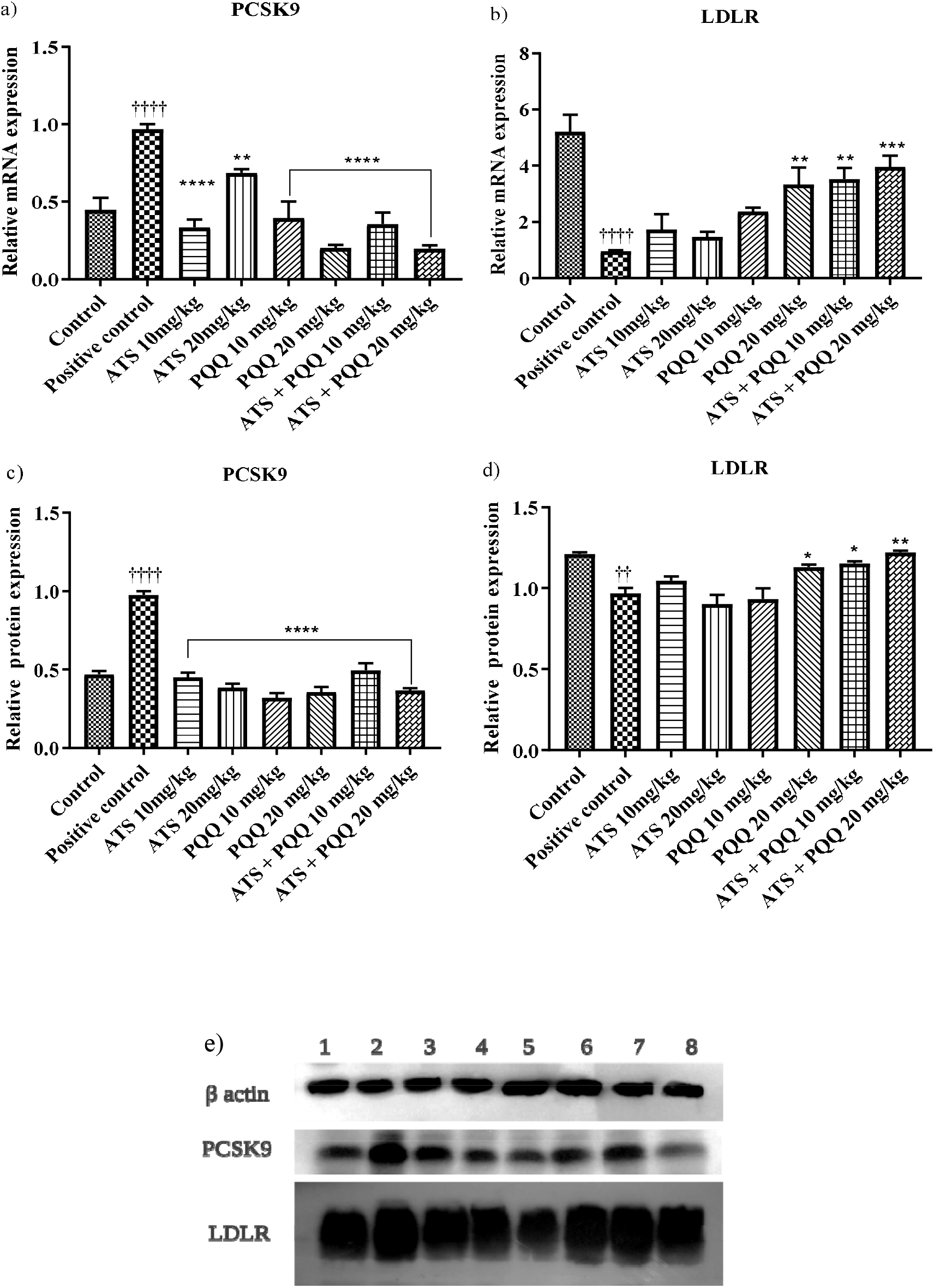
Effect of PQQ and ATS on hepatic mRNA and protein expression. Data are means ± SEM. ^††††^*P* < 0.0001 when compared to the control group. *P* < 0.0001; *P* < 0.001; *P* < 0.01 when compared with positive control group. Statistical significance was determined by one-way ANOVA with Dunnett’s post-hoc test.

Hepatic LDLR protein levels were measured to verify whether PCSK9 inhibition by PQQ decreased serum lipids by preventing LDLR degradation. Hepatic LDLR protein levels were increased after PQQ 20 mg/kg treatment (1.13 ± 0.01, *P* < 0.05) and also along with atorvastatin (1.15 ± 0.01, *P* < 0.05; 1.22 ± 0.01, *P* < 0.01) in the HFFD fed rats compared to positive control. Compared to atorvastatin in the doses of 10 or 20 mg/kg alone (1.04 ± 0.02; 0.9 ± 0.05), the combination treatments distinctly increased LDLR protein levels. This increased fold expression of LDLR is due to remarkable reduction in PCSK9 levels in rats treated with atorvastatin, PQQ alone and its combinations when compared with positive control. Although decrease in PCSK9 concentration was observed in the group treated with atorvastatin *per se* however the ability to restore/recycling of LDLR was not demonstrated as compared to positive control (Figure 3c-e).

## Discussion

Obesity and overweight have been shown to be associated with risk of cardiovascular diseases such as atherosclerosis, myocardial infarction and stroke, which are closely associated with lipoprotein metabolism [26] and increase in chronic low-grade inflammation by macrophage infiltration and enhanced expression of proinflammatory markers [27]. It is well-known that gain in body weight can elevate TG, LDL-C and TC levels and with reduction in HDL-C. In our previous study, the data collected indicated that both atorvastatin and PQQ *per se* and in combination could decrease in chronic low-grade inflammation and augment glucose tolerance levels [17] [20].

In a clinical study by Careskey and colleagues, it was reported that atorvastatin could increase the circulating PCSK9 levels by the activation of transcription factor SREBP-2, that activates both LDLR and PCSK9 genes [28]. Since, PCSK9 is positively associated with TG levels with impact on lipid metabolism [29], the present study was designed to investigate the effects of PQQ alone and in combination with atorvastatin in rats fed with HFFD. PQQ along with atorvastatin has proved decrease in serum TC, TG, LDL-C, VLDL-C, non-HDL and increase in HDL-C as compared to positive control which was also reported in our previous study [20]. The decrease in serum triglycerides, LDL cholesterol and total cholesterol was in accordance with earlier clinical reports.[30] [31]

High atherogenic index and TC/HDL ratio indicates high risk of developing coronary artery disease and non-HDL cholesterol index may be particularly helpful in predicting risk of cardiovascular diseases [32]. Under HFFD exposure, the ratios of TG/HDL, LDL/HDL, CHL/HDL, non-HDL/HDL and atherogenic index, demonstrated higher values, this result was in concurrence with an earlier report [33]. While both atorvastatin, PQQ *per se* and in combination inflicted positive improvement in these clinical end points of the study when compared to positive control. It was noted the outcomes were more pronounced when PQQ was supplemented along with atorvastatin therapy in the rats.

To determine the efficacy of PQQ on HFFD induced hepatocellular injury and related endpoints, we studied histological changes in liver tissues which were published in our earlier reports [20] and we confirmed the histological findings by using biomarkers of liver injury – serum ALT and AST levels which were elevated in HFFD group compared to normal control and were attenuated by the treatment *per se* and PQQ in combination with atorvastatin. Several studies have demonstrated the induction of PCSK9 by statins in cultured cells and in animal models [34] [35] [36]. Our findings in consumption of HFFD eventually led to upregulated PCSK9 and downregulate LDLR when compared with normal control group, is inconsistent with previous reports [37]. These perturbations were reversed and showed significant downregulation of PCSK9 in liver as well as in circulating serum concentrations on treatment with PQQ (10 and 20 mg/kg) *per se* and in combination with ATS when compared to positive control. Treatment with atorvastatin (10 and 20 mg/kg) *per se* demonstrated decreased gene and protein expression of LDLR when compared with normal control group. It also demonstrated no significant change in circulating serum PCSK9 concentration when treated with treatment with atorvastatin (10 and 20 mg/kg) alone as compared to positive control group. Rescue of LDLR from intracellular degradation was verified by an increase in hepatic LDLR protein levels after PQQ treatment and this upregulation of LDLR was further enhanced in combination with atorvastatin. These results support an improvement in cholesterol management by adding PQQ to statin treatment. In human studies, it was reported that atorvastatin at a 40-mg dose significantly elevated circulating PCSK9 protein levels [28].

Extensive clinical trials are being conducted using PCSK9 monoclonal antibodies (mAbs), as an attractive therapy for lowering LDL-C levels [38]. Human mAbs against PCSK9 have been developed to target PCSK9 inhibition, namely, evolocumab and alirocumab. Understanding the complexity of the molecular pathway and its therapeutic strategy, these PCSK9 inhibitors may have unexpected adverse effects resulting from the loss of PCSK9 functions at other sites in the body, in particular regarding neurocognition, muscle toxicity, immunologic and allergic effects and inability to cause plague regression similar to statins, resulting in “legacy effect” after discontinuation [39]. Thus, PQQ in combination with atorvastatin could make PQQ an attractive candidate for enhancing statin efficacy in cost effective way of therapeutic management. Collectively, these was the very first study in demonstrating the role of PQQ on increase expression of LDLR and providing evidence for divergent regulation of PCSK9 transcription of PQQ in combination with atorvastatin.

## Author contributions

KD; conceptualization, investigation, methodology, formal analysis, Writing. DJ; RT-PCR, data curation and formal analysis. RK; RT-PCR, western blot and formal analysis. AM; supervision, conceptualization, formal analysis and corrections.

